# A conserved enhancer cluster regulates *efnb2* expression in the vertebrate dorsal retina

**DOI:** 10.64898/2026.03.30.715377

**Authors:** Lina Shi, Louise Perez, Gabriela Lima, Josane F. Sousa, Zhuoxin Chen, Evgeny Kvon, Igor Schneider, Patricia N. Schneider

**Author notes:** Corresponding author: Patricia N. Schneider., **Email:**.

## Abstract

The vertebrate retina is patterned along its dorsal-ventral (DV) axis by conserved signaling centers, yet the *cis*-regulatory architecture maintaining these territories remains unclear. Using the four-eyed fish *Anableps anableps*, which enables physical separation of dorsal and ventral retina, we integrated RNA-seq, ATAC-seq and Hi-C to map DV regulatory landscapes. We identify a 29-kb dorsal-enriched enhancer cluster within a gene poor region between *efnb2a* and *gtpbp8*, embedded in a conserved topologically associating domain spanning *efnb2a-gtpbp8-slc10a2*. Dorsal-biased chromatin contacts link this cluster to the *efnb2a* promoter. The orthologous human region contains a genome-wide significant association with retinal thickness (rs2575134; P = 2e-14). Leveraging the compact genome of *Tetraodon nigroviridis*, we cloned the orthologous interval and show that it drives dorsal retina-specific reporter expression in zebrafish and mouse. Together, these results identify an evolutionarily conserved regulatory domain controlling retinal axial identity and linking enhancer architecture to human ocular variation. Funding, National Science Foundation.

## Main

Across vertebrates, the retina is patterned along its DV axis into domains with distinct cellular composition and visual functions^1-4^. These territories arise during development through morphogen signaling and regionally expressed transcription factors that establish axial identity and guide retinal circuit organization^5-20^. In particular, Eph-ephrin signaling mediated by ephrin-B2 (*Efnb2*) plays a central role in translating DV positional information into topographic mapping of retinal ganglion cell projections^21,22^.

Although the molecular pathways specifying DV identity have been extensively characterized^23-55^, the *cis*-regulatory mechanisms that control spatial expression of these patterning genes remain poorly understood. Defining these regulatory landscapes requires direct comparison of chromatin accessibility and genome architecture between dorsal and ventral retinal domains. Here we take advantage of the four-eyed fish *Anableps anableps*, whose unusually large eye (Fig. 1a) and expanded ventral retina domain enables physical separation of dorsal and ventral retina, providing an opportunity to interrogate DV-specific regulatory architecture in the adult vertebrate retina^56,57^.

**Fig. 1.**
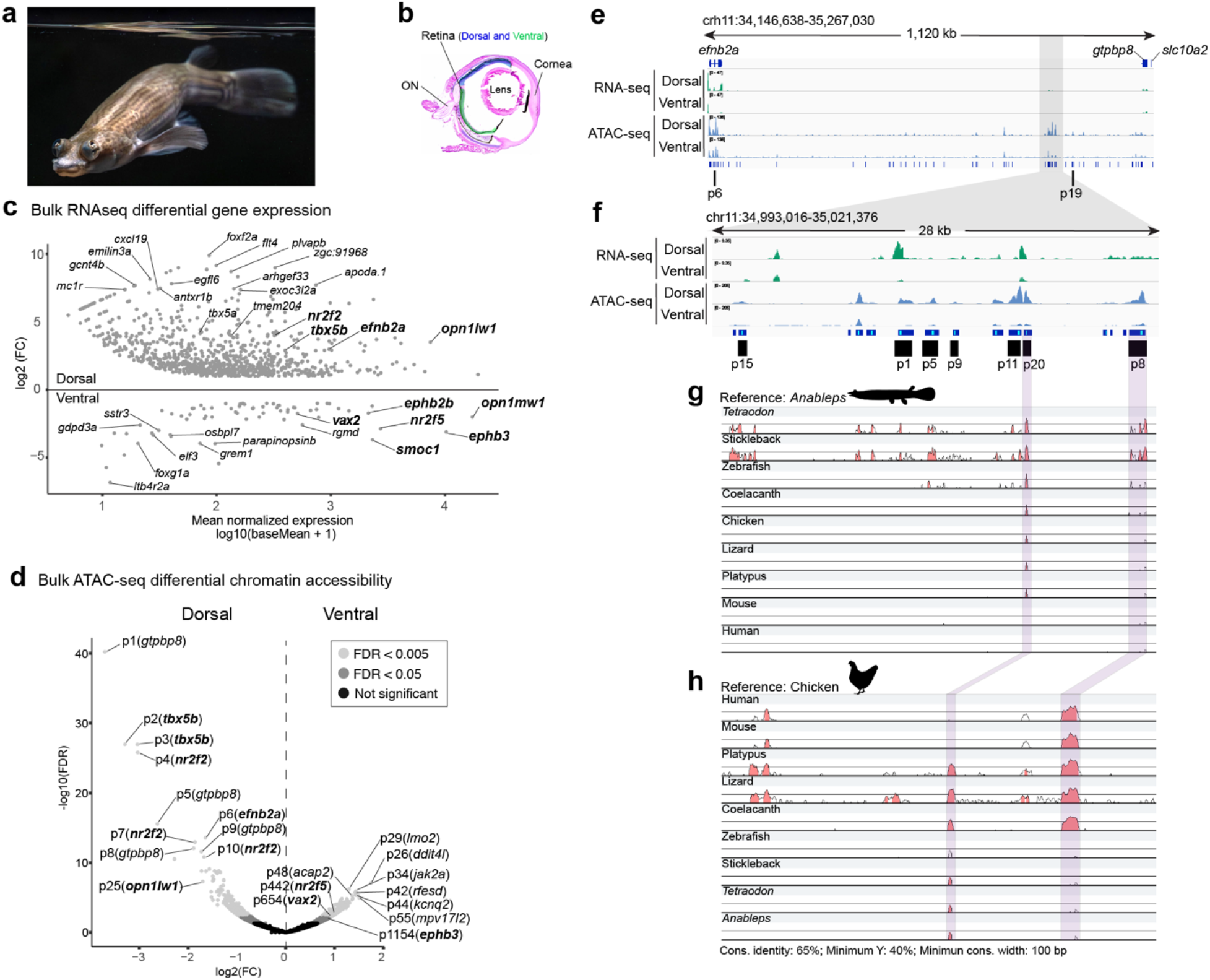
The gene poor region downstream of *efnb2a* harbors a conserved enhancer cluster. **a**, Anableps swimming at the water-air interface. **b**, *Anableps* dorsal (blue) and ventral (green) retina domains. **c**, MA plot showing gene expression differences in the dorsal and ventral retina domains. **d**, Volcano plot showing regions (peaks) of differential chromatin accessibility between dorsal and ventral retina domains and the closest annotated genes near peaks in parenthesis. **e**, RNA-seq and ATAC-seq tracks visualized in the Integrative Genome Viewer (IGV) showing dorsal retina enrichment for efnb2a expression and a cluster of ATAC-seq peaks. **f**, Zoomed in view of IGV tracks in the region of the enhancer cluster showing cluster of dorsal-enriched peaks. **g-h**, mVISTA alignment with *Anableps* (**g**) or chicken (**h**) as references, showing sequence conservation across vertebrates.

In *Anableps*, the ventral retina spans approximately half of the retinal length, as defined by ventral identity markers and ventrally restricted opsin genes^57-59^ (Fig. 1b). To resolve DV regulatory differences, we bisected adult retinas at their approximate midpoint and generated RNA-seq and ATAC-seq profiles from dorsal and ventral hemiretinas. RNA-seq analysis recovered robust DV transcriptional programs, including canonical markers of axial identity. The dorsal retina showed elevated expression of *efnb2a, tbx5b, nr2f2* and *opn1lw1*, whereas the ventral retina preferentially expressed *smoc1, nr2f5, ephb3, vax2* and *opn1mw2* (Fig. 1c; Supplementary Data 1), confirming accurate regional dissection and preservation of DV identity. ATAC-seq profiling identified 245,570 accessible chromatin regions exhibiting DV-biased accessibility. Dorsal-enriched peaks localized near established dorsal regulators (*tbx5b*: p2, p3; *nr2f2*: p4, p7, p10, p14, p18; *efnb2a*: p6) (Fig. 1d; Supplementary Data 2), while ventral-enriched peaks were detected near *vax2* (p654), *nr2f5* (p442) and *ephb3* (p1154), consistent with transcriptional polarity.

Notably, a striking concentration of highly significant dorsal-accessible peaks mapped to a gene-poor interval between *efnb2a* and *gtpbp8* (p1, p5, p8, p9, p11, p15, p19, p20), highlighting this locus as a candidate regulatory hotspot (Fig. 1e; Supplementary Data 2). Given the density and magnitude of differential accessibility within this interval, we examined the region in greater detail. We identified a discrete cluster of seven strongly dorsal-enriched peaks (p1, p5, p8, p9, p11, p15, p20) spanning ∼29 kb and coinciding with elevated dorsal *efnb2a* expression (Fig. 1f). Comparative genomic analysis using mVISTA revealed conservation of multiple elements within this cluster across vertebrates. Using either *Anableps* (Fig. 1g) or chicken (Fig. 1h) as reference genomes, we detected conserved sequences corresponding particularly to peaks p20 and p8, retained from teleosts to amniotes. Together, these results identify a dorsal retina-specific enhancer cluster embedded within a conserved genomic interval adjacent to *efnb2a*, defining a candidate *cis*-regulatory module underlying DV retinal identity.

The identification of a dorsal-enriched enhancer cluster within a gene-poor interval adjacent to the dorsally expressed *efnb2a* locus suggested distal regulatory control of *efnb2a*. To define three-dimensional chromatin architecture in this region, we generated high-resolution Hi-C maps from bisected dorsal and ventral adult retina. In the dorsal retina, Hi-C revealed a well-defined chromatin interaction domain spanning ∼2.5 Mb on chromosome 11 (Fig. 2a). This domain encompasses *efnb2a* at its 5′ boundary and extends toward *gtpbp8* and *slc10a2* at its 3′ boundary, forming a structurally coherent TAD. Within this domain, we identified a prominent looping interaction (Loop 41) that connects the *efnb2a* promoter to the distal enhancer cluster located upstream of *gtpbp8* and *slc10a2*. In addition to Loop 41, the *efnb2a* promoter engages in multiple contacts across the same distal interval, indicating a highly interactive regulatory hub. Integration with ATAC-seq confirmed that the loop-engaged distal region corresponds to a dense cluster of chromatin-accessible elements that are strongly enriched in the dorsal retina and markedly reduced in the ventral retina (Fig. 2a). Quantitative comparison of loop strengths between retinal domains showed that Loop 41 is among the most significantly dorsal-biased interactions (FDR = 1.9e-4; Fig. 2b; Supplementary Data 3), positioning it as a dominant dorsal-specific chromatin contact linking the enhancer cluster to *efnb2a*.

**Fig. 2.**
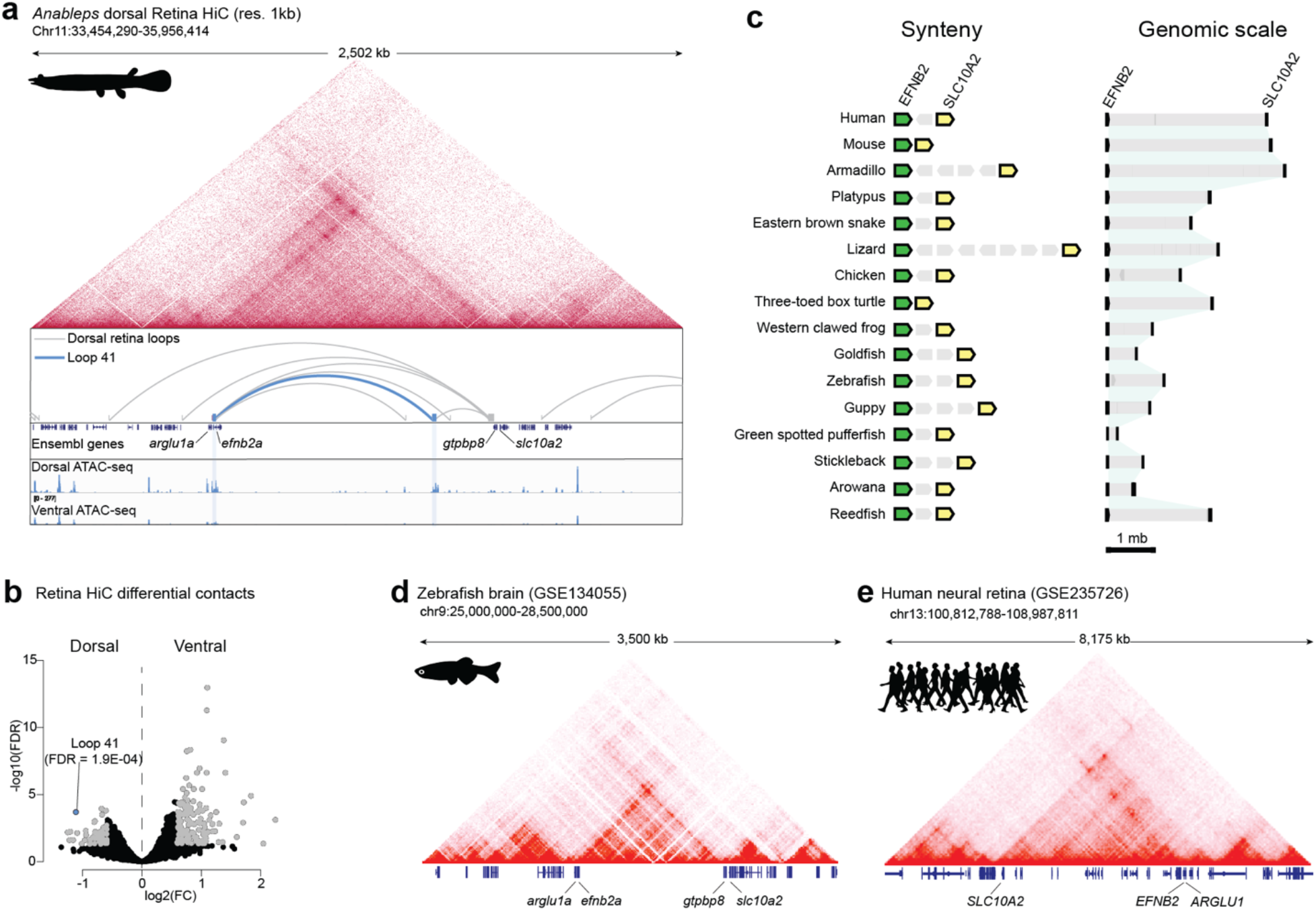
Hi-C identifies dorsal-enriched loop connecting the enhancer cluster to efnb2a. **a**, *Anableps* Hi-C data showing TAD structure at the *enfb2*-*slc10a2* genomic interval; Loops show contacts occurring within the TAD; loop 41 (blue) links the enhancer cluster with *efnb2a* gene; ATAC-seq tracks show dorsal-enriched peaks in the enhancer cluster and in *efnb2a*. **b**, Volcano plot showing differential contacts between dorsal and ventral retinal; loop 41 ranks among the most significant dorsal-enriched contacts. **c**, Analysis of synteny and gene distances shows conservation of a large gene-poor region encompassing *EFNB2* and *SLC10A2* genes across vertebrates. **d-e**, Zebrafish (**d**) and human (**e**) Hi-C datasets confirm evolutionary conservation of TAD structure spanning *EFNB2* and *SLC10A2* genes.

To assess evolutionary conservation of this regulatory architecture, we examined genomic organization across 16 vertebrate species. Synteny analysis revealed consistent linkage of *EFNB2* and *SLC10A2* across mammals, reptiles, amphibians and teleosts (Fig. 2c), with preservation of an extended intergenic interval often spanning megabases. The orthologous interval resides within a TAD in both zebrafish (Fig. 2d) and human (Fig. 2e), mirroring the structural organization observed in *Anableps*. This conservation supports a model in which long-range enhancer-promoter communication within a structurally constrained topological domain represents an ancient regulatory strategy. Together, these findings demonstrate that a dorsal-enriched enhancer cluster physically engages the *efnb2a* promoter through a conserved long-range chromatin loop embedded within a stable TAD, revealing domain-level regulation of DV retinal identity.

Given the deep conservation of the *efnb2-slc10a2* intergenic interval, we asked whether the orthologous human region harbors genetic variation associated with retinal phenotypes. Query of the GWAS catalog identified eighteen variants, including rs2575134, a tri-allelic intergenic variant significantly associated with retinal thickness (P = 2e-14) in 43,148 individuals of European ancestry^60^ (Supplementary Data 4). Interval-wide inspection revealed that among variants within the broader conserved locus, rs2575134 represents the strongest retina-associated signal. Alignment of the human locus with the *Anableps* enhancer cluster using mVISTA demonstrated that rs2575134 resides within the orthologous conserved interval upstream of EFNB2 and next to the region corresponding to *Anableps* peak 8 (Fig. 3a). Genome browser annotation revealed that the variant falls within an ENCODE candidate *cis*-regulatory element (cCRE) and lies adjacent to regions marked by H3K27ac in multiple cell types, consistent with regulatory activity in this interval (Fig. 3b). The surrounding locus exhibits a dense distribution of predicted *cis*-regulatory elements, reinforcing its characterization as a regulatory hotspot. Higher-resolution inspection of the variant-containing sequence revealed predicted transcription factor binding motifs, including a RORB motif positioned immediately adjacent to rs2575134 (Fig. 3c). RORB is a nuclear receptor with established roles in retinal development and photoreceptor differentiation^61,62^, raising the possibility that allelic variation at rs2575134 may influence transcriptional regulation. The localization of rs2575134 within a conserved enhancer-rich interval orthologous to the dorsal-enriched enhancer cluster identified in *Anableps* suggests that this region may contribute to modulation of EFNB2 expression and human retinal thickness variation.

**Fig. 3.**
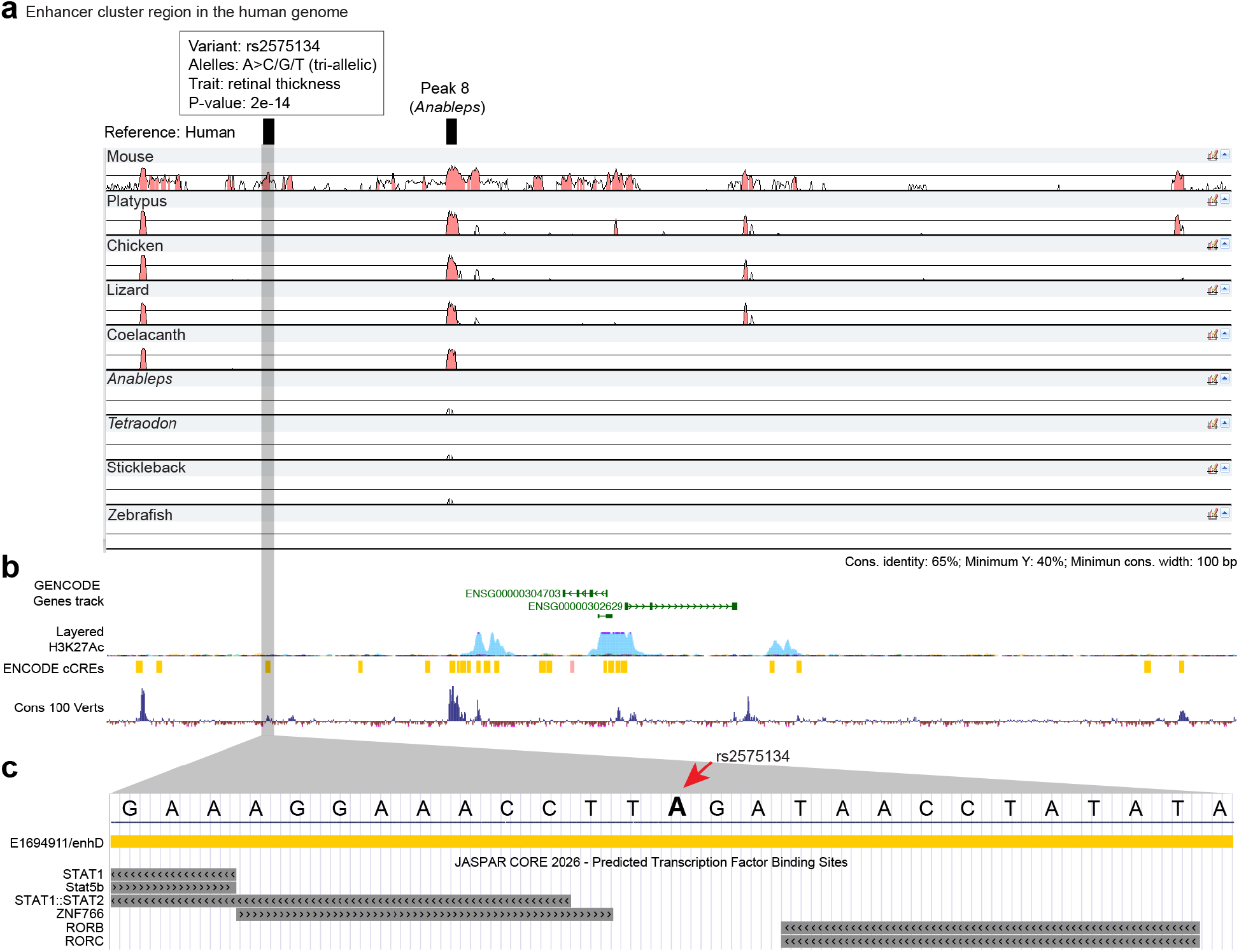
The human orthologous enhancer cluster region harbors a genetic variant associated with retinal thickness. **a**, mVISTA alignment with human as reference showing the orthologous location of the conserved *Anableps* enhancer cluster peak 8 and the location of variant rs2575134, associated with retinal thickness. **b**, Genome Browser tracks showing the presence of non-coding genes, H3K27Ac marks and ENCODE cCREs within this region; variant rs2575134 mapped to E1694911/enhD. **c**, Zoomed in view of the variant rs2575134 location showing its position relative to predicted transcription factor binding sites.

The enhancer cluster identified in *Anableps* spans nearly 30 kb, posing a practical challenge for functional dissection. To overcome this limitation, we leveraged the compact genome of the green spotted pufferfish *Tetraodon nigroviridis* (Fig. 4a), in which orthologous regulatory intervals are often substantially reduced in size^63^. We first established whether DV retinal domains in *Tetraodon* exhibit transcriptional and regulatory profiles comparable to those observed in *Anableps*. Because clear morphological landmarks demarcating DV retinal boundaries are absent in *Tetraodon*, we estimated the ventral domain to span approximately the ventral third of the retina, as observed in other fully aquatic teleosts^64^. Retinas were therefore dissected into dorsal and ventral fractions corresponding to approximately two-thirds and one-third of the retinal length along the DV axis (Fig. 4b). RNA-seq profiling recovered canonical DV transcriptional programs consistent with those observed in *Anableps*. Dorsal markers including *efnb2a, nr2f2*, and *tbx5b* were enriched in dorsal retina samples, whereas *smoc1, ephb2*, and *vax2* showed higher expression in ventral retina (Fig. 4c; Supplementary Data 5). Cross-species comparison confirmed strong concordance in expression domain, fold change, and overall expression levels of conserved DV markers between *Anableps* and *Tetraodon* (Fig. 4d). ATAC-seq profiling similarly revealed DV-biased chromatin accessibility patterns. Dorsal-enriched peaks mapped near key dorsal regulators including *efnb2a* and *nr2f2*, and near *gtpbp8*, while ventral-enriched peaks near *vax2* and *smoc1* were uncovered (Fig. 4e; Supplementary Data 6). Notably, peaks adjacent to *gtpbp8* ranked among the most strongly dorsal-enriched sites, mirroring the regulatory architecture observed in *Anableps*.

**Fig. 4.**
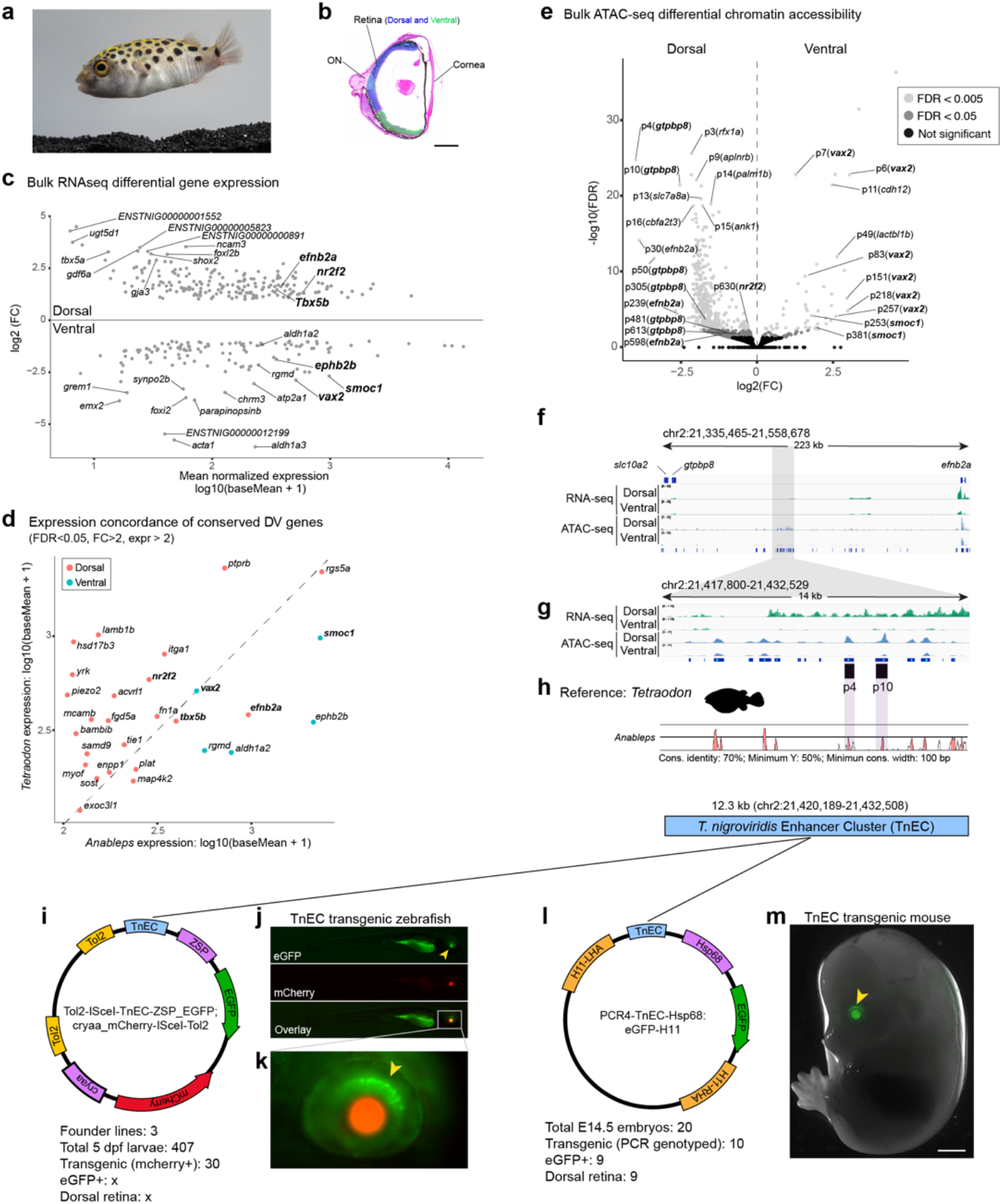
The compact Tetraodon enhancer cluster drives dorsal retina reporter gene expression in transgenic zebrafish and mouse. **a-b**, *Tetraodon* (**a**) and its estimated dorsal (blue) and ventral (green) retina domains (**b**). **c**, MA plot showing gene expression differences in the dorsal and ventral *Tetraodon* retina domains. **d**, Concordant expression typical dorsal and ventral markers in *Anableps* and *Tetraodon*. **e**, Volcano plot showing regions (peaks) of differential chromatin accessibility between dorsal and ventral retina domains and the closest annotated genes near peaks in parenthesis. **f**, RNA-seq and ATAC-seq tracks visualized in IGV showing dorsal retina enrichment for *efnb2a* expression and a cluster of ATAC-seq peaks. **g**, Zoomed in view of IGV tracks in the region of the enhancer cluster showing cluster of dorsal-enriched peaks. **h**, mVISTA alignment with *Tetraodon* as reference showing extensive sequence conservation overlapping with dorsal-enriched ATAC-seq peaks; diagram of the orthologous *Tetraodon* enhancer cluster region cloned for heterologous transgenesis assays. **i**, Diagram of transgenesis construct used to generate stable zebrafish lines. **j**, Transgenic zebrafish (5 dpf); eGFP, mCherry and overlay are shown. **k**, Zoomed in region eGFP expression in the dorsal retina. **l**, diagram of transgenesis construct used to generate mouse zebrafish lines. **m**, transgenic mouse embryo (E.14.5) showing eGFP expression in the dorsal retina. Arrowheads (yellow) show eGFP expression in the lens and retina of the zebrafish (**j**,**k**) and mouse (**m**).

Inspection of RNA-seq and ATAC-seq tracks across the genomic interval spanning *efnb2a* and *gtpbp8* revealed that dorsal-enriched peaks p4 and p10 reside within a compact cluster of accessible elements selectively enriched in dorsal retina (Fig. 4f,g). Comparative sequence alignment using mVISTA confirmed conservation of these elements between *Tetraodon* and *Anableps* (Fig. 4h), supporting the interpretation that this region represents the orthologous enhancer cluster identified in *Anableps*.

Consistent with the extensive genome compaction characteristic of pufferfishes, the putative *Tetraodon* enhancer cluster (TnEC) spanned approximately 14 kb, or half the size of the corresponding interval in *Anableps*. This compact organization enabled cloning of a 12.3-kb fragment encompassing the cluster of dorsal-accessible peaks for functional assays.

To test regulatory activity of this interval, we leveraged well-established transgenic reporter constructs in zebrafish (Fig. 4i) and mouse (Fig. 4l). F1 transgenic progeny from three independent zebrafish lines carrying the TnEC reporter consistently displayed GFP expression in in the dorsal region of the developing lens (Fig. 4j) and retina (Fig. 4k). Remarkably, transgenic mouse embryos harboring the same interval driving GFP expression also exhibited reporter activity in the dorsal retina and lens (Fig. 4m). The dorsally restricted retinal activity of TnEC across these distantly related vertebrates, together with the phylogenetic distances separating teleosts and mammals (∼450 million years), demonstrates deep functional conservation of this regulatory module. These findings provide functional evidence that the enhancer cluster identified near *efnb2* constitutes an evolutionarily conserved regulatory element associated with dorsal retinal identity.

Retinal DV patterning is established early during development through morphogen gradients and transcription factors, yet the *cis*-regulatory landscape that promotes these patterning cues has remained poorly defined. Here we identify a cluster of dorsal-enriched enhancers located in a gene-poor interval adjacent to *efnb2a*, a key component of the Eph-ephrin signaling system that mediates DV retinal mapping. Integration of chromatin accessibility, gene expression, and Hi-C data in *Anableps* reveals that this enhancer cluster resides within a large TAD and forms dorsal-biased chromatin contacts with the *efnb2a* promoter. These findings provide a regulatory framework linking classical DV patterning genes to a spatially restricted *cis*-regulatory module.

Several lines of evidence indicate that this regulatory architecture is deeply conserved across vertebrate evolution. The *efnb2*-*slc10a2* genomic neighborhood is conserved across diverse vertebrate lineages, and the orthologous interval resides within a comparable TAD in zebrafish and human genomes. In *Anableps*, within this TAD, multiple enhancer elements show sequence conservation, some retained from teleosts to amniotes, suggesting long-term preservation of regulatory function. At the same time, not all enhancer elements within the cluster display detectable sequence conservation. Recent comparative chromatin studies have proposed that retinal gene regulation is governed by conserved *cis*-regulatory codes that persist across vertebrates despite turnover of individual enhancer sequences^65^. Our findings are consistent with this view and suggest that the broader regulatory landscape surrounding *efnb2a*, including the clustering of enhancer elements and their organization within a stable chromatin domain, has been maintained across vertebrate evolution even as individual enhancer units evolve.

Notably, the high density of accessible elements, evidence of non-coding transcription, and association with a key developmental regulator are features reminiscent of enhancer clusters described as super-enhancers that control genes governing cell identity and developmental patterning^66,67^. Similar clustered regulatory architectures have been described at other retinal loci, including the modular super-enhancer controlling *VSX2* expression during retinal development^50^. The compact genome of *Tetraodon* allowed us to isolate an orthologous interval encompassing this enhancer cluster and test its activity experimentally. Strikingly, this interval was sufficient to drive dorsal retina-restricted reporter expression in both zebrafish and mouse, species separated by nearly 450 million years of evolution. Together, these results indicate that the regulatory logic controlling dorsal *efnb2a* expression is embedded within an ancient enhancer cluster whose function has been maintained across vertebrates.

Our findings also suggest that this conserved regulatory interval may contribute to variation in human retinal architecture. A genome-wide significant association with retinal thickness maps within the orthologous human interval, within a predicted *cis*-regulatory element. Although the precise regulatory targets of this variant remain to be established, its position within a dense regulatory landscape corresponding to the enhancer cluster described here raises the possibility that genetic variation in this domain modulates retinal structure through effects on *EFNB2*. More broadly, our study illustrates how combining spatially resolved genomics, comparative genomics, and cross-species functional assays can reveal deeply conserved regulatory domains underlying vertebrate tissue patterning, providing a framework for linking developmental gene regulation with evolutionary conservation and human phenotypic variation.

## Methods

### Animal work

*Anableps, Tetraodon* and zebrafish were maintained and used in accordance with approved Louisiana State University (LSU) IACUC protocols IACUCAM-25-094 and IACUCAM-24-003. Fish were obtained from commercial vendors. *Anableps* and *Tetraodon* were maintained in large tanks with a recirculating freshwater system at 27-28 °C under a 12 h light/12 h dark cycle. Zebrafish were kept in small tanks within a recirculating freshwater system at 27-28 °C and a 12 h light/12 h dark cycle. Prior to eye dissection, animals were euthanized in 0.3 % MS-222. Mice were maintained under standard housing conditions and timed pregnancies were established as described previously^68^ and in accordance with approved University of California Irvine IACUC protocol numbers AUP-20-001 and AUP-23-005. Embryos were collected at E14.5, and genotyping was performed following the published protocol^68^.

### Tissue flash-frozen, cryosection and histology

Dissected eyes from adult *Anableps* and Tetraodon were flash-frozen using isopentane cooled with liquid nitrogen. Isopentane was dispensed into a metal container and placed in liquid nitrogen to equilibrate for about 10 minutes. Tissues were embedded in Tissue-Tek O.C.T. compound (Sakura) using appropriate molds for 30 minutes. Using forceps, the molds were placed into the equilibrated isopentane until frozen. Samples were stored at -80 °C for sectioning. The tissues were cryosectioned at 10 µm (thickness) at -20 °C using a Leica CM1520 cryostat. And the slides were stored at -80 °C until further use. Hematoxylin and Eosin (H&E) staining was performed as previously described^69^.

### RNA isolation, library preparation, and sequencing and data analysis

Total RNA was extracted from individual TRIzol (Life Technologies) preparations via a phenol-chloroform extraction. Retinal tissue was first dissociated by vigorous pipetting in TRIzol, following manufacture’s instructions (Thermo Scientific). RNA concentration and purity were assessed with a BioDrop (Thermo Scientific). For transcriptome sequencing, four biological replicates were prepared for dorsal and ventral retina from *Anableps*, and three biological replicates were prepared for dorsal and ventral retina from *Tetraodon*. In each case, 500 ng of total RNA per sample was used for library preparation with the NEBNext Ultra II RNA Library Prep Kit (Illumina), following the manufacturer’s instructions. Paired-end sequencing was performed on an Illumina NovaSeq 6000 platform. The reference genome for *Anableps* (GCA_014839685.1) and Tetraodon (GCA_051020865.1) together with corresponding gene annotation files were downloaded from the National Center for Biotechnology Information (NCBI) webpage. All RNA sequencing runs were mapped to reference genomes using STAR^70^. After mapping, we used featureCounts (Subread package) to count the number of reads mapped to each gene^71^. Differential gene expression (DGE) analysis between the dorsal and ventral retinas was measured using DESeq2^72^.

DESeq2 utilizes a model based on the normalization of reads by the median of ratios^73^. Before running DESeq2, we performed pre-filtering to retain only genes with more than 10 reads. P-values were adjusted using Benjamini and Hochberg’s method to control the false discovery rate (FDR). Genes with adjusted p-values < 0.05 and a log2 fold change greater than 1 or less than -1 were considered differentially expressed. RStudio was used to create principal component analysis (PCA) and MA plots.

### Assay for Transposase-Accessible Chromatin using sequencing (ATAC-seq)

ATAC-seq was performed on freshly dissociated nuclei from bisected dorsal and ventral retinas of *Anableps* (4 individuals) and Tetraodon (three individuals). Tagmentation and library preparation were carried out using the Active Motif ATAC-seq Kit (Cat. No. 53150, Carlsbad, CA, USA) following manufacturer’s protocol. Briefly, dissociated nuclei were quantified using a hemocytometer, and approximately 50,000 nuclei were incubated with Tagmentation Master Mix (25 µl 2x tagmentation buffer, 2 µl 10x PBS, 0.5 µl 1.0% Digitonin, 0.5 µl 10% Tween 20, 12 µl nuclease-free water, 10 µl Assembled Transposomes) in a thermomixer with 800 rpm agitation. The tagmented DNA was purified and amplified by PCR. The resulting libraries were sequenced pair-end using the NovaSeq 6000 platform (Novogene Corporation Inc., Sacramento, CA, USA) generating 75 million paired-end reads of 150 bp (NCBI Sequence Read Archive project number XX). Adapter sequences were removed from the reads using NGmerge (version 0.3)^74^. The trimmed reads were aligned to the *Anableps* and Tetraodon reference genomes using Bowtie2 (version 2.2.5)^75^. Alignments were output as a BAM file, which was indexed using Samtools (version 1.18)^76^. The fragment size distribution was assessed using the ATACseqQC package (version 1.24.0)^77^ in RStudio (version 4.3.1). The trimmed reads were name-sorted using Samtools (version 1.18)^76^ and further processed with Genrich (version 0.6.1) to call narrow peaks using the parameters -j -y -r -v -m -e -a -f, to remove duplicate reads, multimapped reads, and reads mapped to mitochondrial chromosomes. Downstream analysis of the ATAC-seq data was performed using DiffBind (version 3.10.1) to identify differential chromatin accessibility between dorsal and ventral retina domains (peaks)^78^. Peak sets generated by Genrich were associated with metadata for each sample and imported into DiffBind within RStudio (version 4.3.1). Read counts were generated using 75 summits, and the reads were normalized using DESeq2 (version 1.40.2)^72^. Peaks that exhibited differential accessibility between the retina domains were selected based on a significance threshold of FDR < 0.05. The differential peaks between the dorsal and ventral retina were visualized using the Integrative Genomics Viewer (IGV) (version 2.14.1). Subsequently, we used the script annotatePeaks.pl from HOMER v5.1^79^ to identify and annotate genes located near the peaks.

### Hi-C Library Preparation, sequencing and analysis

Hi-C was performed on freshly bisected dorsal and ventral retina tissue of *Anableps* (n = 4; 2 adult males and 2 adult females) using the Arima-Hi-C+ Kit (Arima Genomics, Cat. No. A510008, Carlsbad, CA, USA), following the manufacturer’s protocol. Briefly, tissues were crosslinked with formaldehyde and digested with a restriction enzyme cocktail provided in the kit. Proximity ligation was performed to join spatially proximal DNA fragments, followed by purification and fragmentation of the ligated DNA. Biotinylated fragments were enriched and processed for library preparation using the Arima Library Prep Module (Arima Genomics, Cat. No. A303011, Carlsbad, CA, USA). Libraries were sequenced on the NovaSeq 6000 (Novogene Corporation Inc., Sacramento, CA, USA) using 150bp paired-end reads, yielding approximately 600 million read pairs per sample. Hi-C analysis was performed using HOMER v5.1^79^ with an Arima-specific in silico-digested *Anableps* genome. Adapters were removed using the homerTools trim script (5’ trimming), and reads were aligned independently to the *Anableps* genome using Bowtie2 v 2.2.5^75^. After alignment, we generated a paired-end tag directory for each replicate using makeTagDirectory using the parameters -tbp 1 and -removePEbg. Loops and TADs were identified using findTADsAndLoops.pl with a 3 kb resolution and an FDR threshold of 0.05. All loops and TADs were merged per condition using merge2Dbed, after which we quantified loops and TADs across all replicates and conditions. Differentially enriched loops and TADs were identified using getDiffExpression.pl with simpleNorm and edgeR parameters. Finally, we generated the hic file using tagDir2hicFile.pl with Juicer auto parameters. The loops were visualized in the Integrative Genomics viewer v2.14.1 software and the TADs in the genome browser website. Differential contacts were visualized via Volcano plot generated in RStudio (version 4.3.1). Hi-C datasets and images from zebrafish brain (GSE134055) and human neural retina (GSE235726) were downloaded from the 3D Genome Browser 2.0 website^80^.

### Synteny and genomic architecture

The gene organization and genomic distances from 16 vertebrate species corresponding to the gene interval spanning EFNB2 and SLC10A2 were obtained and adapted from Genomicus v106.01^81^ website.

### Cloning of constructs

To create the construct for zebrafish transgenesis, the *Tetraodon* enhancer cluster sequence was cloned into Tol2-ISceI-ZSP_EGFP; cryaa_mCherry-IsceI-Tol2 vector^82^ using NheI digestion followed by Gibson assembly (NEB, E2621). For mouse transgenesis, the *Tetraodon* enhancer cluster sequence was cloned into the PCR4-Hsp68::eGFP-H11 vector^68^ (Addgene #211941) using AgeI digestion followed by Gibson assembly (NEB, E2621). The *Tetraodon* enhancer cluster sequence was obtained via PCR amplification from *Tetraodon* genomic DNA extracted using the Quick-DNA Miniprep Plus Kit (ZYMO, cat. No. D4069). PCR was performed using either Q5 High-Fidelity 2X Master Mix (NEB, M0492) or repliQa HiFi ToughMix (Quantabio, Part No. 95200). For both constructs in this study, whole-plasmid sequencing (Eurofins Genomics) was performed to ensure the integrity of the vector and enhancer cluster sequences before injection. Primers used for cloning (with backbone sequences underlined) were as follows: TOL2-Forward: 5′ [sequence] 3′ and TOL2-reverse: 5′ [sequence] 3′, PCR4-Forward: 5′ [TCTTCAGGCTGAAGCTGATGG AACAGGTACCCTCATTTGCTTTGGCTGGCG] 3′ and PCR4-reverse: 5’ [CGGCTGCTCAGTTTGGATGTTCCTGGCGGCCGCCAGGGTTCATCTGCTGGTTT] 3′.

### Imaging

Images were acquired using a Nikon Eclipse Ni fluorescent microscope with a 2x, 10x or 20x lenses. Brightness and contrast were adjusted uniformly across entire images for clarity in figure presentation. Raw, unprocessed image files are available for all datasets. H&E slides were visualized and imaged using a Nikon Eclipse Ni microscope with a Nikon DS-Fi3 camera.

### Transgenic mouse generation

Transgenic mice in this study were generated using a CRISPR/Cas9 microinjection protocol as previously described^68^. Briefly, a mix of Cas9 protein (20 ng/μl; IDT, Cat. No. 1074181), sgRNA (50 ng/μl), and circular donor plasmids (7 ng/μl) in injection buffer (10 mM Tris, pH 7.5; 0.1 mM EDTA) was injected into the pronucleus of FVB/NJ embryos. Donor plasmids were column-purified using a PCR purification kit (Qiagen) and eluted in injection buffer prior to injection. Superovulated female FVB/NJ mice (7-8 weeks old) were mated to FVB/NJ stud males, and fertilized embryos were collected from oviducts. F0 embryos were either brought to gestation or collected at E14.5.

### Zebrafish transgenic lines

A freshly prepared solution containing 100 ng/μl of the transgenesis vector, 100 ng/μl of transposase mRNA, and a trace amount of Fast Green for visualization was microinjected into one-cell stage zebrafish (Danio rerio) embryos. Injected embryos were maintained in E3 medium supplemented with PTU at 28.5 °C and screened at 5 days post-fertilization (dpf) for mosaic GFP and mCherry fluorescence. F0 embryos showing the highest levels of mosaic GFP:mCherry expression were selected for imaging.

### mVista alignment

Genomic sequences spanning the intergenic region between the *efnb2a* and *gtpbp8*/*slc10a2* genes were extracted from eight species and submitted to the mVISTA web server (http://genome.lbl.gov/vista/mvista/)^83^ using default settings to identify conserved sequences. The *Anableps* Anableps, Tetraodon, or chicken genomes were used as reference sequences for alignments. Sequence alignments were performed with the Shuffle-LAGAN algorithm, and conservation profiles were generated using a 100 bp sliding window with identity set at 65% or 70% and a minimum conservation threshold of 40% or 50% sequence identity. Conserved non-coding regions identified within the enhancer cluster region were extracted for downstream analyses of potential regulatory function.

### Genome-wide association (GWAS) variant identification within the EFNB2 regulatory interval

To identify genetic variants associated with ocular phenotypes within the conserved regulatory interval surrounding EFNB2, we queried the genomic region chr13:103,888,078-104,080,831 (GRCh38), corresponding to the orthologous enhancer cluster identified in this study. Variants and associated GWAS signals within this interval were retrieved from the NHGRI-EBI GWAS Catalog using the gwasrapidd R package. Queries were performed using genomic coordinates spanning the full interval. This analysis identified eighteen variants, with rs2575134 (chr13:103,996,957) as the only variant within the interval associated with a retinal phenotype. rs2575134 is an intergenic variant associated with retinal thickness (GWAS Catalog study GCST90528011; P = 2e-14) in a cohort of 43,148 individuals of European ancestry^60^. Association metadata including risk allele and study details were obtained directly from the GWAS Catalog

### Cross-species expression concordance analysis

Differential expression tables were imported into RStudio (version 4.3.1) and processed using dplyr, readxl, and ggplot2. Genes were retained if they met the following thresholds in each species: adjusted P value < 0.05, absolute fold change > 2, and expression level ≥ log10(baseMean + 1) of 2. Fold-changes were oriented such that positive values corresponded to dorsal enrichment. Gene identifiers were converted to lowercase and duplicates within each species were collapsed by retaining the entry with the lowest adjusted P value and highest expression. Orthologous genes were considered concordant when the same gene symbol was detected in both species and exhibited the same DV direction of differential expression. Expression concordance was visualized by plotting log10(baseMean + 1) expression values for the matched gene set between species.

## Data availability

Raw sequencing data for bulk RNA-seq, ATAC-seq, and Hi-C seq generated in this study are deposited in the Gene Expression Omnibus (GEO) under the accession numbers: GSE310896 (RNA-seq of *Anableps*), GSE311047 (RNA-seq of *Tetraodon*), GSE310894 (ATAC-seq of *Anableps*), GSE310895 (ATAC-seq of *Tetraodon*), and GSE322638 (Hi-C of *Anableps*). All code used in this study is available at our GitHub repository: https://github.com/lnperez90/Shi-et-al-2026.

## Acknowledgments

We thank Emily Kane and Joe Fontenot for animal photography. This work was funded by start-up funds from Louisiana State University (P.S. and I.S), and an NSF-Integrative Organismal Systems (IOS) grant (2409931, P.S.).

## Contributions

Conceived the study: I.S. and P.S. Study design: L.S., I.S., and P.S. Data generation and quality control analyses: L.S., L.P., G.L., J.F.S., Z.C., E.K., I.S. and P.S. Analysis and interpretation: L.S., L.P., G.L., J.F.S., Z.C., E.K., I.S. and P.S. Wrote the initial draft: L.S., I.S., and P.S. Read and provided comments on the manuscript: L.P., G.L., J.F.S., Z.C., E.K. Supervised the project: P.S.

